# Do tigers hunt during the day? Diel Activity of the Asian tiger mosquito, *Aedes albopictus* (Diptera: Culicidae), in Urban and Suburban Habitats of North America

**DOI:** 10.1101/2021.05.05.442733

**Authors:** Isik Unlu, Ary Faraji, Nicholas Indelicato, James R. McNelly

## Abstract

*Aedes* (*Stegomyia*) *albopictus* (Skuse) impacts human outdoor activity because of its aggressive biting behavior, and as a major vector of mosquito-borne diseases, it is also of public health importance. Although most mosquito species exhibit crepuscular activity by primarily host seeking at dawn and dusk, *Ae. albopictus* has been traditionally characterized as a diurnal or day-biting mosquito. With the global expansion and increased involvement of *Ae. albopictus* in mosquito-borne diseases, it is imperative to elucidate the diel activity of this species, particularly in newly invaded areas. Human sweep netting and carbon dioxide-baited rotator traps were used to evaluate the diel activity of *Ae. albopictus* in two study sites. Both trapping methods were used in New Jersey’s Mercer County, USA (temperate urban), while only human sweep netting was used in Florida’s Volusia County, USA (subtropical suburban). Human sweep netting was performed to determine adult mosquito activity at sunrise, solar noon, sunset, and lunar midnight. Because New Jersey is in a temperate area, diel activity was investigated during the early season (3-19 July), peak season (25 July-19 September), and late season (22 September-22 October). *Aedes albopictus* showed the highest activity during peak and late seasons at solar noon (*P* < 0.05). At sunrise and sunset during the peak season, *Ae. albopictus* activity was similar. Lunar midnight activity was significantly lower than sunrise and solar noon (*P* < 0.05) but was similar to that of sunset. In the late season, the highest activity was observed during solar noon while the least activity was observed during sunrise and lunar midnight (*P*<0.05). Rotator traps used in conjunction with the human sweep net technique exhibited similar results. Seasonal activity was not differentiated in Florida due to the consistent subtropical weather. The highest adult activity was observed at sunrise using human sweep netting but it was not significantly different from solar noon and sunset. The lowest adult activity was observed at lunar midnight; however, it was not significantly different from solar noon and sunset. These results provide evidence that the diel activity of *Ae. albopictus*, contrary to the common perception of its diurnal activity, is much more varied. Because of the involvement of the species in the transmission of debilitating mosquito-borne pathogens such as chikungunya, dengue, and Zika virus, coupled with its affinity to thrive in human peridomestic environments, our findings have global implications in areas where *Ae. albopictus* thrives. It also highlights the importance of behavioral studies of vector species which will not only help mosquito control professionals plan the timing of their control efforts but also provide empirical evidence against conventional wisdoms that may unjustly persist within public health stewards.

**Author Summary:** The Asian tiger mosquito, *Aedes albopictus*, is an invasive mosquito which is now established in at least 40 states in the USA. Lack of efficient surveillance and control methods against *Ae. albopictus*, in addition to human-aided accidental transportations, have played a great role in its rapid expansion. Although surveillance measures are becoming more systematic and effective, control of this species still poses a great challenge. *Aedes albopictus* is difficult to control in the larval stage because it primarily develops in artificial containers that are widespread in peridomestic habitats. These habitats are not only ubiquitous in these environments, they are also cryptic, inaccessible, and extremely difficult to control. Therefore, control of *Ae. albopictus* in these environments often relies on adult control measures which utilize insecticides dispersed through ultra-low volume equipment as a cold aerosol space spray. These adulticide applications are often conducted at night against endemic mosquito species which are primarily active between dawn and dusk. However, since *Ae. albopictus* has been traditionally classified as a day-biting mosquito, mosquito control specialists have had doubts about the efficacy of a nocturnal application against a diurnally active mosquito. These uncertainties about intervention efforts become even more important during public health outbreaks of mosquito-borne pathogens such as chikungunya, dengue, or Zika viruses when protection of public health is of paramount importance in peridomestic habitats. Our investigations provide evidence that *Ae. albopictus* exhibits activity throughout the day and night and that nighttime adulticide applications may indeed be effective against this species, and should not be disregarded.

## INTRODUCTION

Most terrestrial organisms are exposed to daily changes in light, dark, and temperature cycles. They have adapted to these changes and express specific behaviors which are genetically controlled. *Aedes albopictus* (Skuse), the Asian tiger mosquito, is a competent vector of many mosquito-borne viruses such as dengue (DENV) and chikungunya (CHIKV) [1,2]. It is also a major pest species that can drive children indoors and detrimentally impact human quality of life [3,4]. Understanding the diel activity of *Ae. albopictus*, specifically the times when it may be host seeking, is essential because of its vectorial status as well as the need for effective control measures.

The need for successful and sustainable *Ae. albopictus* control programs became more evident due to the recent outbreak of arboviral diseases globally, particularly with the expansion of Zika virus (ZIKV) [5]. Additionally, outbreaks of DENV in locations such as the Seychelles Islands, China, La Réunion, Hawaii, Mauritius, and Europe have implicated *Ae. albopictus* as the primary vector [6,7]. Autochthonous transmission of CHIKV implicating *Ae. albopictus* as the main vector, has also been documented both in France [8] and Italy [9]. In Gabon, central Africa, epidemiological surveillance has determined that *Ae. albopictus* was the principal vector of ZIKV during an urban outbreak in 2007 [10]. The species has also drawn the attention of vector control and public health professionals, particularly in expanding and newly invaded areas [11–13].

When an invasive species becomes endemic in a new area, it may display different biological behaviors, including host preference, diel activity, and vector competence [10,14–16], which all pose new challenges for vector control specialists. The first detection of *Ae. albopictus* in the continental United States in 1985 and its subsequent spread displayed that mosquito control districts were not equipped with effective methods to conduct surveillance and control for this peridomestic species. For example, the establishment of *Ae. albopictus* in New Jersey was first recorded in 1995 from a trap collection in Keyport, Monmouth County [17], however it wasn’t until 2008 that all 21 mosquito control districts in New Jersey finally obtained an effective trap to conduct surveillance for this species [18,19]. Prior to this, the presence of *Ae. albopictus* was only detected using New Jersey light traps, which are poor devices in gauging the presence or abundance of this species [20]. In the late 2000’s, after numerous field evaluations, the newly created Biogents Sentinel (BGS) (Biogents AG, Regensburg, Germany) traps were recognized as the standard surveillance tool for *Ae. albopictus* and *Aedes aegypti* L. [20–22]. Even though the BGS traps are generally operated over a 24-hour period, they do not provide information about the diel host seeking periods of invasive *Aedes* mosquitoes such as *Ae. aegypti* and *Ae. albopictus.* Elucidating the diel activity of these species is crucial to understanding their behavior, which helps vector control and public health professionals better focus their surveillance and control efforts to maintain quality of life and prevent disease outbreaks.

In Hawley’s (1988) review of *Ae. albopictus* biology, field studies on diel activity patterns conducted in several countries in Asia reported peak blood feeding during daylight but rarely during the night hours [23]. Almeida et al. (2005) reported from the Chinese Territory of Macao, that *Ae. albopictus* displayed a bimodal biting peak activity during the morning and later afternoon [24]. Similar results were reported on La Reunión Island where bimodal blood feeding activity for *Ae. albopictus* females was higher in the morning and afternoon peaks [25]. Hassan (1996) found morning and evening twilight peaks for both sexes of *Ae. albopictus* in Malaysia [26]. On the contrary in Japan, researchers determined active *Ae. albopictus* behavior through night time nectar feeding between 2100 and 2130 hours [27]. In addition, although researchers observed bimodal activity in Macao, they also detected some activity during all 24 hours of the day for *Ae. albopictus* [24]. However, the vast majority of previously published investigations all incriminate *Ae. albopictus* as primarily diurnal [24–26,28]. This has led to the acceptance of certain fallacies that have permeated vector control communities, particularly in the USA. For example, adult mosquito suppression methods generally utilize adulticides which are applied as ultra-low volume (ULV) cold aerosol sprays during the night. But because ULV applications have not been efficacious or long lasting in controlling diurnally active urban mosquitoes, they have been declared ineffective, particularly for reduction of disease transmission, as reviewed in [29]. One reason for failure of control has been attributed to the nocturnal resting behavior of day-biting mosquitoes in natural and artificial places that are sheltered from the insecticide plume. The ineffectiveness of nighttime ULV applications against diurnal mosquitoes has unfortunately become the conventional wisdom within the modern vector control community and many vector control programs simply do not attempt to adulticide against *Ae. albopictus* because they are under the assumption that this species may not be active at all during the nighttime ULV application periods.

However, during a variety of field investigations aimed at elucidating the biology, ecology, and effective control methods against *Ae. albopictus* in temperate central New Jersey [30–32], it has been observed that this species may also be active even during the nighttime hours [29,33]. As a result, the goal of this study was to investigate the diel activity of *Ae. albopictus* in New Jersey and Florida, in order to further elucidate the biology of this important vector mosquito. Our primary objective was to provide empirical data to prove that invasive *Aedes* mosquitoes, such as *Ae. albopictus*, are indeed active throughout the 24 hr diel period and to challenge conventional wisdoms that night time applications of adulticides may indeed be effective during those periods because of the continuous activity of the target species around the clock.

## Materials and Methods

### Site Selection

All study sites in New Jersey were highly urbanized, residential sites within the City of Trenton and were comparable to the field site descriptions provided by Unlu et al. (2011) and Farajollahi et al. (2012). The study area encompassed a mix of two-story row homes or duplexes and occasionally, abandoned homes subject to occupation by transients. Study sites in Florida were selected in suburban neighborhoods in the City of Edgewater. Edgewater is located along the Indian River, adjacent to the Mosquito Lagoon.

### Human Sweep Net Collections

Human sweep net collections were made using a standard 30.5 cm diameter sweep net purchased from Fisher Scientific (Atlanta, GA, USA). In both locations, the same individuals conducted human sweep netting for the duration of the experiment. With two collectors per residential properties, the collectors took turns walking around the perimeter of the parcel, as the geography allowed, and ending at the pre-determined sampling location which was a shaded or partially shaded area free of obstruction. At the sampling location, each collector stood still and moved the sweep net in a figure eight pattern (Fig. 1) for five minutes to collect mosquitoes. On each minute mark, the collector rotated 90° degrees, such that in the final minute the individual was in the original (first minute) position. After completing the sampling process, the first collector returned to the vehicle to place the net in a cooler with dry ice and after 15 minutes, the second collector repeated the process with a second net. A single collector performed all sampling in Florida following the same experimental protocol used in New Jersey. All mosquito specimens collected were counted and identified to species. Weekly collections were made between 8 August and 22 October in New Jersey and between 16 July and 9 September 2013 in Florida. For analysis of New Jersey data, the study was divided into two seasonal periods: 1) peak (8 - 29 August), and 2) late (5 September - 3 October).

**Fig. 1.**
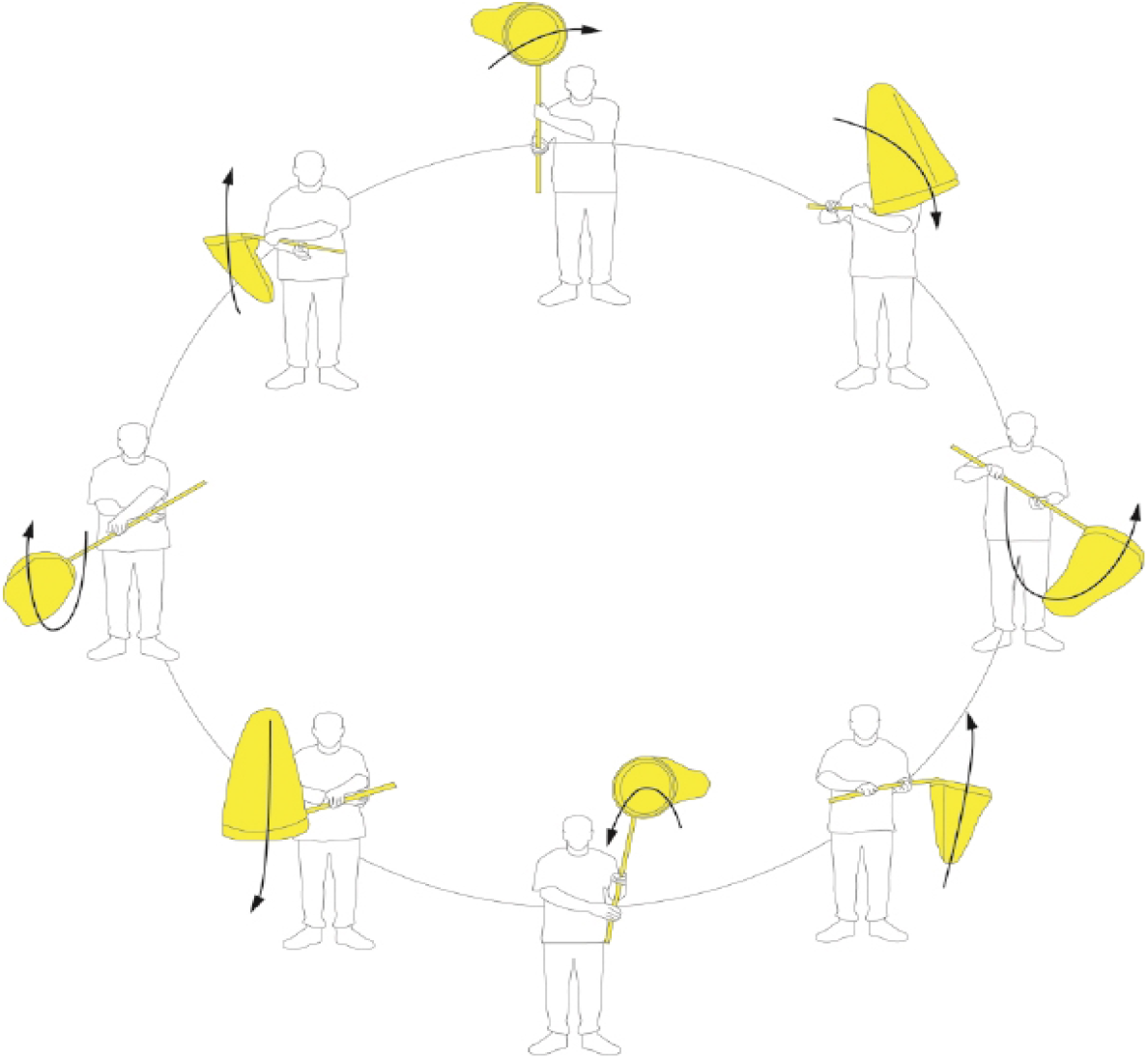
Diagramatic representation of individual performing human sweep net surveillance using a butterfly or “Figure 8” pattern.

### Sampling Time – Diel Periods

A 24 h day was divided into four discrete sampling periods: Sunrise, Solar Noon, Sunset, and Lunar Midnight. Solar Noon was identified using the National Oceanic & Atmospheric Administration solar calculator (http://www.esrl.noaa.gov/gmd/grad/solcalc/) and Lunar Midnight was defined as 12 h post Solar Noon. Sunrise and Sunset were delineated using Weather Underground’s regional times for sunrise and sunset (https://www.wunderground.com/). Ten and six residential properties were chosen in New Jersey and Florida, respectively, which required that the human sweep netting be initiated 30 mins prior and ending no more than 30 mins post the determined sampling periods.

### Carbon Dioxide-baited Bottle Rotator Trap Collection

In New Jersey the diel activity of *Ae. albopictus* was also measured using a collection bottle rotator (CBR) equipped with a CDC miniature light trap (model 1512 and 512 respectively, John W. Hock Company, Gainesville, FL, USA). The CBR consists of a programmable timer powered by a 12-volt, ten-amp rechargeable DC battery which allows for the collection of adult mosquitoes at eight different times during a 24 h period. A voltage regulator (BioQuip Products Inc., Ranco Dominguez, CA, USA) was added to allow the use of the 6-volt CDC miniature trap on the 12-volt CBR system. A CBR with eight jars was used to segregate collections into eight time periods, each lasting three hours. The periods were defined as: 0700-1000, 1000-1300, 1300-1600, 1600-1900, 1900-2200, 2200-0100, 0100-0400, and 0400-0700. Traps were re-programmed weekly to compensate for seasonal changes according to Solar Noon. The CBR traps were baited with CO_2_ and BG Lures (Biogents AG, Regensburg, Germany). Traps were held in place using a cast-concrete base and positioned 0.5 m above ground level. Adult mosquito collections took place between 3 July and 22 October 2013. At the end of 24 h sampling period, traps and trap contents were transported to the laboratory, identified to species, and counted. Female and male *Ae. albopictus* were recorded separately. For data analysis, the study was divided into three seasonal periods: 1) early (3-19 July), 2), peak (25 July-19 September), and 3) late (22 September-22 October).

### Comparison of Biogents Sentinel Trap Catches with Human Sweep Net Collection

Biogents Sentinel traps were deployed in conjunction with human sweep net surveillance in order to investigate the correlation between the two. Biogents Sentinel traps were baited with a BG-Lure, and deployed for a 24 h sampling period. Traps were deployed in the same parcels where human sweep netting was performed. The traps were placed no closer than 7.5 m and no more than 10.5 m away from the location of the human sweep netting. At the end of 24 hours, traps and trap contents were transported to the respective laboratories in New Jersey and Florida, identified to species, and counted. Female and male *Ae. albopictus* were recorded separately.

#### Data Analysis

##### Analysis of HSN Data

To determine the peak activity for *Ae. albopictus*, we compared the number of adults, female and male, collected during four time periods using a generalized linear model [34]. Overdispersion was detected via the Poisson model (value/df = 1.986), and the model was refit using negative binomial distribution (PROC GENMOD, SAS version 9.3; SAS Institute 2011) with site, season, time and interaction term season*time all used as predictors. The model was determined to fit the data adequately (χ^2^ = 76.62, df = 63, *P* = 0.116) in New Jersey and the interaction term was significant (*χ^2^* = 65.20, df = 3, *P* < 0.001). The association between Florida human sweep netting counts and time were investigated using Poisson regression adjusted for overdispersion (PROC GENMOD, SAS version 9.3; SAS Institute 2011), with site and time as predictors. The model fit adequately (*χ*^*2*^ = 20.61, df =15, *P* = 0.150) and the main effect of time was significant ((*χ*^*2*^ = 9.94, df = 3, P = 0.020). The *P* values between comparisons were adjusted using Holm’s test, which adjusts the calculation of probability in line with the number of comparisons made to avoid type I errors (Holm 1979).

##### Collection Bottle Rotator Trap Analysis

In order to compare the peak activity for *Ae. albopictus* using both human sweep netting and CBR trapping, two consecutive time periods from the CBR trapping collections were combined as follows: Sunrise collection was comprised of 05:00-11:00 collections; 11:00-17:00 collections formed Solar Noon; Sunset was comprised of 17:00-23:00; and 23:00-05:00 collections formed Lunar Midnight. The analysis was performed using the generalized linear model (PROC GENMOD, SAS version 9.3; SAS Institute 2011). Overdispersion was detected in Poisson and negative binomial models. Therefore, an ANOVA model was fit to the bottle rotator trap data. The association between season and time were investigated using ANOVA (PROC GLM, SAS version 9.3 for Windows). Since normality and equal variances assumptions were violated, the Box-Cox transformation [35] was used to achieve approximate normality and the eighth root transformation was used to normalize the data. The main effects of site, season and time were used as predictors of the transformed adult counts. The *P* values between comparisons were adjusted by using Tukey’s tests.

##### Comparison Between HSN and BGS Trap Collections

Linear correlations (Pearson’s correlation) were calculated between the numbers of *Ae. albopictus* collected for human sweep netting and BGS traps for each state. Each human sweep netting session for each trapping location was 10 min while BGS traps deployed for 24 hrs, therefore a linear correlation test was appropriate to investigate the concordance of the two sampling methods.

## RESULTS

### Human Sweep Net

A total of 808 *Ae. albopictus* adults were collected in New Jersey, with 374 (46.3%), specimens collected during Solar Noon followed by 172 (21.3%) at Sunrise, and 156 (19.3%) and 106 (13.1%) during Sunset and Lunar Midnight, respectively. Of the total number captured, 508 were females (62.9%) along with 300 males (37.1%). The association between human sweep netting counts, season, and time were investigated with site, season, time, and season*time as predictors. The main effect of season*time was significantly associated with the collections (χ^2^ = 65.20, df = 3, *P* < 0.001). All pairwise contrasts for season*time were examined and the results are listed in Table 1. Human sweep netting collections showed the greatest activity during Solar Noon in New Jersey during the peak and late season. Substantial activity was also detected at Sunrise, Sunset, and Lunar Midnight during the peak season with no significant difference between Sunset and Lunar Midnight. The mean number of *Ae. albopictus* per collection declined in late season (Figure 2). The highest level of activity was at Solar Noon, followed by that of Sunset, and both were statistically different (*P* < 0.05) from Sunrise and Lunar Midnight.

**Table 1.** Least square means from log-linear analysis of New Jersey human sweep netting counts of all *Aedes albopictus* for ten study sites over two seasons (peak and late) and four time periods: Sunrise, Solar Noon, Sunset, and Lunar Midnight.

| Season | Time | LS Mean<sup>‡</sup> | 95% C.I. |
| --- | --- | --- | --- |
| Peak | Sunrise | 13.48<sup>b</sup> | 10.60 - 17.15 |
|  | Solar Noon | 22.73<sup>a</sup> | 18.30 - 28.23 |
|  | Sunset | 8.07<sup>c</sup> | 6.12 - 10.64 |
|  | Lunar Mid | 8.77<sup>c</sup> | 6.70 - 11.5 |
| Late | Sunrise | 0.39<sup>c</sup> | 0.14 - 1.05 |
|  | Solar Noon | 10.22<sup>a</sup> | 7.89 - 13.25 |
|  | Sunset | 3.62<sup>b</sup> | 2.51 - 5.23 |
|  | Lunar Mid | 0.10<sup>c</sup> | 0.01 - 0.70 |

<sup>‡</sup>Values not sharing the same superscript are significantly different (Holm's test, $P < 0.05$ )

**Fig 2.**
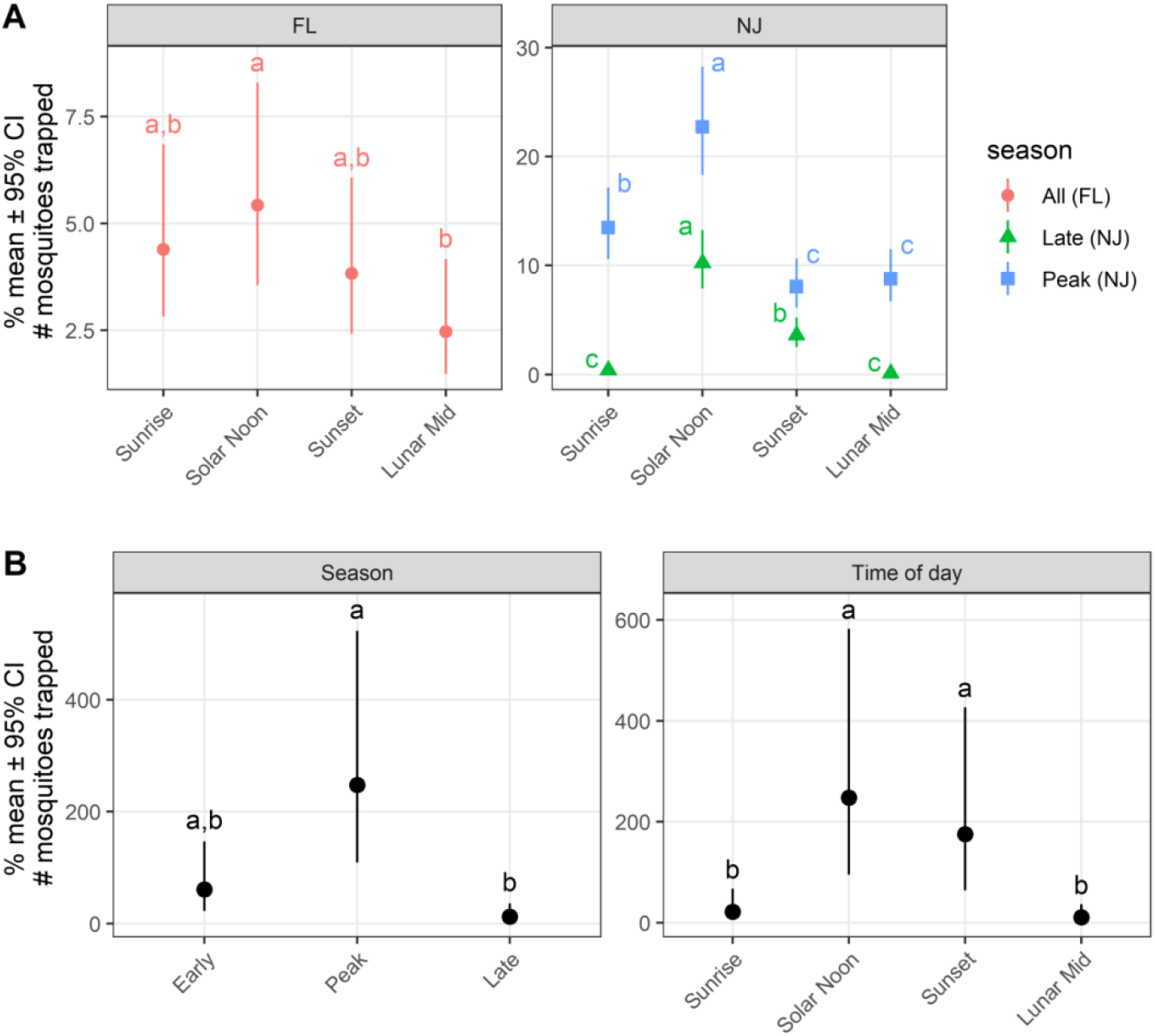
A. Human sweep netting counts from Florida for all *Aedes albopictus* during four time periods; Sunrise, Solar Noon, Sunset, and Lunar Midnight (red bars), and human sweep netting counts from New Jersey for all *Aedes albopictus* over two seasons (peak and late) and four time periods: Sunrise, Solar Noon, Sunset, and Lunar Midnight. B. Count bottle rotator trap counts of all *Aedes albopictus* at two study sites over three seasons (early, peak, and late) and over a 24 hr period in New Jersey. ^¥^ Values not sharing the same superscript are significantly different (Holm’s test, P < 0.05)

In Florida, a total of 202 *Ae. albopictus* specimens were sampled with 68 (33.7%) captured at Sunrise, followed by 55 (27.2%) at Solar Noon, 48 (23.8%) during Sunset and 31 (15.4%) at Lunar Midnight. Collections were weighted towards females with a ratio of 5:1. The main effect of time was significantly associated with the collections (χ^2^ = 9.94, df=3, *P* = 0.020). All pairwise contrasts for time were examined and the results are listed in Table 2. The only statistically significant activity for *Ae. albopictus* adults were found during Solar Noon and Lunar Midnight (Figure 2).

**Table 2.** Least square mean estimates from log linear analysis of Florida human sweep net counts of all *Aedes albopictus* for six sites over four time periods: Sunrise, Solar Noon, Sunset, and Lunar Midnight.

| Time | LS Mean<sup>‡</sup> | 95% C.I. |
| --- | --- | --- |
| Sunrise | 4.39<sup>a,b</sup> | 2.82 - 6.85 |
| Solar Noon | 5.43<sup>a</sup> | 3.56 - 8.29 |
| Sunset | 3.83<sup>a,b</sup> | 2.42 - 6.07 |
| Lunar<br>Midnight | 2.47<sup>b</sup> | 1.47 - 4.17 |

<sup>‡</sup>Values not sharing the same superscript are significantly different (Holm's test, $\alpha < 0.05$ )

### Collection Bottle Rotator Trap

We tried several statistical models for analysis of the CBR trap data set. We found the best fit with ANOVA for CBR trap collections. The main effects of site (F(1,17) = 13.79, *p*= 0.002), season (F(2,17)=11.40, *p* = 0.001), and time (F(3,17) = 9.03, *p* = 0.001) were significant predictors of the transformed mosquito counts. The least squares means and corresponding confidence intervals along with the results of post-hoc Tukey’s tests for season and time are provided in the Tables 3 and 4. *Aedes albopictus* adult activity was highest during peak season followed by early and late season (Figure 2). For time periods, the highest activity was recorded during Solar Noon and it was significantly different than the other times investigated, excluding Sunset (Table 4). The second highest activity was recorded during Sunset and it was significantly different than Sunrise and Lunar Midnight.

**Table 3.** Least square means with confidence intervals for New Jersey bottle rotator trap counts of all *Aedes albopictus* at two study sites over three seasons (early, peak, and late) and over a 24 hr period.

| Season | LS Mean | 95% C.I. |
| --- | --- | --- |
| early | 61.08<sup>a,b</sup> | 22.68 - 146.98 |
| peak | 247.92<sup>a</sup> | 109.26 - 522.93 |
| late | 12.48<sup>b</sup> | 3.66 - 35.96 |

Values not sharing the same superscript are significantly different (Tukey's test, $P < 0.05$ )

**Table 4.**
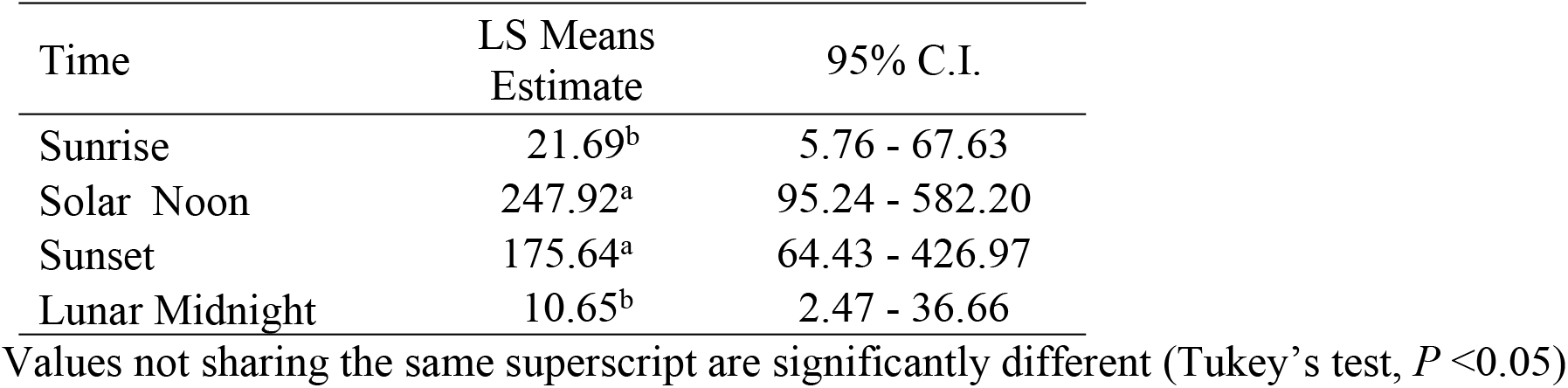
Least square means with confidence intervals for New Jersey bottle rotator trap counts of all *Aedes albopictus* at two study sites over three seasons (early, peak and late) combined, during four different time periods.

| Time | LS Means<br>Estimate | 95% C.I. |
| --- | --- | --- |
| Sunrise | 21.69<sup>b</sup> | 5.76 - 67.63 |
| Solar Noon | 247.92<sup>a</sup> | 95.24 - 582.20 |
| Sunset | 175.64<sup>a</sup> | 64.43 - 426.97 |
| Lunar Midnight | 10.65<sup>b</sup> | 2.47 - 36.66 |

Values not sharing the same superscript are significantly different (Tukey's test, $P < 0.05$ )

### Comparison between HSN and BGS Trap Collections

A total of 808 *Ae. albopictus* adults were collected in New Jersey during human sweep netting collections. The BGS traps collected a total of 1,061 adults, close to a 1:1 ratio (585 females and 476 males). In Florida, a total of 202 *Ae. albopictus* were collected with human sweep netting and BGS traps captured a total of 349 adults with a 6:1 ratio (299 females and 50 males). Correlation analysis of mosquito collections showed that HSN and BGS trap collections were positively correlated in New Jersey (r = 0.477, *p* < 0.001) and in Florida (r = 0.401, p < 0.001).

## DISCUSSION

In New Jersey, diurnal activity was clearly greatest at Solar Noon during the peak and late seasons, but did not differ significantly between the two seasons. Activity was different when comparing Sunrise and Sunset in both peak and late season. Interestingly, activity at Lunar Midnight during peak season was only different from than that of Solar Noon. Sampling activity in Florida was initiated in mid-July, when *Ae. albopictus* populations are often high. Overall, *Ae. albopictus* activity levels were lower in the suburban Florida environment than those found in New Jersey’s urban habitat. There were no statistically significant differences in diel activity between Sunrise, Solar Noon, or Sunset in Florida, and while *Ae. albopictus* activity was lowest at Lunar Midnight, the level of activity at this time period was only different from that of activity at Sunrise.

Bimodal activity has been documented in a variety of *Aedes* species including *Ae. aegypti* in Trinidad [36], *Aedes polynesiensis* Marks [37] in Samoa, and *Aedes woodi* Edwards [38] in East Africa. The bimodal activity of *Ae. albopictus* reported by Hawley (1988) and others, including Delatte et al. (2010) on the island of La Réunion during an outbreak of CHIKV was not observed in either New Jersey or Florida [23,25]. Delatte et al. (2010), however, recorded continuous activity across 24h in a series of human-baited experiments [25]. This is also the case with the data generated by human sweep netting collections in New Jersey and Florida, USA. These findings corroborate the previous observations made in New Jersey [29,33] supporting earlier field operations of nocturnal host seeking activity, and research including that of Yee and Foster (1992), Higa et al. (2000), and Barnard et al. (2011), identifying nocturnal host-seeking by *Ae. albopictus* under both laboratory and field conditions [39–41]. Barnard et al. (2011) found that 25% of all *Ae. albopictus* activity monitored by human landing rates took place at night [40]. For those organizations charged with the deployment of ULV space sprays, our research provides supporting evidence for the success of adulticiding directed at *Ae. albopictus* in Mercer between 2009-2011 [29], where nighttime applications of a pyrethroid insecticide (DUET™ Clarke, Roselle, IL, USA) at mid-label rates, when sprayed and spaced one or two days apart, achieved over 80% reduction in *Ae. albopictus* populations in the same urban habitats in which our current studies took place.

This study demonstrated that, late season diel activity was reduced from that of the peak season in temperate New Jersey climate. Activity levels remained greatest at Solar Noon while activity at Sunrise and Lunar Midnight were negligible. Seasonal influences upon the diel activity of *Ae. albopictus* have been identified on La Reunion, with diel activity at night being reduced, and during winter being entirely absent as a result of lower temperatures [25]. Decreasing temperatures in New Jersey are known to influence diel activity of *Aedes* species, and cause shifting levels of activity between the summer and fall seasons [42]. While a temporal shift in the activity of *Ae. albopictus* is not apparent in New Jersey, overall activity is curtailed while diel activity remained limited to warmer temperatures between Solar Noon and Sunset.

Differences in the diel periodicity of *Aedes* mosquitoes from urban and rural locations have also been reported previously [43], and it has been proposed that the increased lumens associated with street and residential lights promotes modifications or adaptations in behavior, including host-seeking. Kawada et al. (2005) performed cross sections of the compound eye of *Ae. albopictus* to determine ommatidial diameter and interommatidial angle, determinants of vision [44]. The eye parameter value (1.59) identified explains the lower level of light that initiated host-seeking in *Ae. albopictus* when compared to *Ae. aegypti* in a laboratory setting, and may explain why *Ae. albopictus* has a preference for brighter environments. In addition, Sippell and Brown (1953) reported the importance of movement to host location by diurnal species [23,45]. For example, under laboratory conditions, similar sized moving targets attracted twice the number of diurnally active *Aedes* mosquitoes compared to stationary targets. Trap capture of *Ae. albopictus* in Japan was attributed to the movement of field personnel walking toward and attending traps [46]. While our research was not designed to determine the potential influence of artificial light (i.e. street lights) in the urban and suburban residences, these might have influenced *Ae. albopictus* night time activity [43]. Similar to artificial light, use of human sweep netting for surveillance must have provided strong visual cues to host-seeking mosquitoes. Biogents Sentinel traps have proven to be an effective surveillance tool for monitoring host-seeking populations of *Ae. albopictus*, and are used routinely to estimate population levels and direct decisions on control activities for this species. It is common practice in Florida to use a landing rate in association with a homeowner’s service request in order to determine if the request is generated by *Ae. albopictus* activity [47]. Biogents Sentinel traps have previously been determined to approximate human landing rate estimates [21,48]. The positive correlation of BGS traps with human sweep netting in both the urban and suburban environments evaluated during this study is invaluable, when considering landing rate surveillance and the influence of varied levels of attractiveness, and collection and enumerating skills when performed by different individuals [49]. In addition, the availability of a surveillance tool such as the BGS trap, removes any perceived ethical concerns or problems related to personnel performing landing rates during periods of disease activity [50].

The diel activity demonstrated by *Ae. albopictus* in both New Jersey and Florida increases the potential of mosquito-human contact and therefore places individuals at risk of health impacts related to arbovirus transmission. The ongoing global expansion of *Ae. albopictus* and dynamics of viral adaptation and vector evolution continues to place more humans at risk [6,13,51]. The introduction of exotic pathogens such CHIKV, DENV, and ZIKV are no longer merely conjectural, but are now considered as regular ongoing occurrences globally. Additionally, DENV is considered the second most important mosquito-borne pathogen, trailing only malaria, which primarily affects impoverished populations and can be considered as a neglected tropical disease. Given the emergence and re-emergence of DENV in both developed and undeveloped regions, and the lack of resources available for effective and timely intervention efforts, our study elucidates additional biological behaviors and diel activity that may prove instrumental for focused control. Furthermore, understanding temporal activity and potential seasonal influences upon levels of activity are also vitally important to the deployment of successful control strategies [52–54]. For personnel actively engaged in organized mosquito surveillance and control programs, this research provides critical information that supports the potential to impact adult populations of a likely disease vector outside widely accepted parameters. These types of investigations will further elucidate the biology and behavior of important vector species, and ultimately lead to more rapid and efficacious intervention efforts against neglected tropical diseases. Our investigations definitively provide evidence that *Ae. albopictus* displays some level of activity throughout the entire day, and that adult mosquito control measures conducted at night should not be discounted as part of an effective integrated vector management program.

## ACKNOWLEDGEMENTS

The authors thank Charles S. Apperson for his critical review of an earlier draft of this manuscript. We thank James Pulaski for creation of drawings and Garret W Dow for field assistance. This work was supported by the USDA National Institute of Food and Agriculture Hatch project accession number 1020755 through the New Jersey Agricultural Experiment Station, Hatch project NJ08530.

